# Evidence of *Escherichia coli* Regulating *Drosophila melanogaster* Behavior

**DOI:** 10.1101/2025.08.20.671414

**Authors:** Hazem Al Darwish, Tia Hart, Deep Patel, Sammi Russo, Safiyah Salama, Muqaddasa Tariq, Jennifer S Sun

## Abstract

Chemosensory systems are critical for insect survival, enabling host seeking, food acquisition, and oviposition site selection. While insect-associated microbes are known to influence host development and immunity, their role in modulating chemosensory behavior remains poorly understood. Here, we demonstrate that *Escherichia coli*, a bacterium identified in non-gut tissues of *Aedes albopictus* and experimentally reintroduced into axenic *Drosophila melanogaster*, alters both larval and adult sensory-driven behaviors. In larvae, *E. coli* infection modified phototaxis and mechanosensory responses across genotypes, while tunneling and thermosensory behaviors were specifically dependent on the ionotropic receptor co-receptor IR25a. In adults, *E. coli* increased attraction to fermentation cues (apple cider vinegar, ethanol) and enhanced sucrose consumption in wild-type and *Orco*-deficient flies, but not in *IR25a*-deficient mutants. Gas chromatography–mass spectrometry revealed that *E. coli* shifted cuticular hydrocarbon composition toward shorter-chain alkanes and increased the sex pheromone 9-tricosene in an IR25a-dependent manner. Together, these findings show that *E. coli* broadly reprograms insect sensory behavior, with IR25a serving as a critical mediator of microbial influence on chemosensory and physiological traits. This work identifies a previously unrecognized role for *E. coli* in shaping insect behavior and chemical ecology, providing a foundation for investigating microbial contributions to host–microbe coevolution and for exploring microbial cues as novel, environmentally sustainable tools for insect control.

**Importance:** Insects depend on their sense of smell and taste to find food, mates, and safe places to reproduce. These behaviors drive the spread of insect-borne diseases and cause major agricultural losses worldwide. Our work shows that a common bacterium, *Escherichia coli*, can reshape insect sensory behaviors, influencing how flies respond to light, temperature, food, and even the chemical signals on their outer surface that control mating. Importantly, these effects are linked to a specific sensory receptor, IR25a, which is conserved across many insect species. This finding reveals a previously unknown role for microbes in fine-tuning insect behavior. Understanding how bacteria influence insect sensory systems not only provides insight into host–microbe interactions but also opens the door to new, environmentally friendly ways of managing insect pests by targeting their behavior rather than killing them outright.

## Introduction

Insects are among the most abundant and diverse animals on Earth, and their success is tightly linked to chemosensory systems that guide feeding, reproduction, and oviposition (1–4). Perturbations to these sensory pathways reduce fitness, foraging efficiency, and survival. With insecticide resistance increasing and chemical controls posing ecological risks (5, 6), identifying endogenous and exogenous regulators of insect chemosensation is essential for developing sustainable control strategies.

Microbes are emerging as critical modulators of insect physiology, influencing immunity, metabolism, and behavior (7, 8). Endosymbionts such as *Wolbachia* alter olfactory receptor expression and odorant-binding proteins, thereby shaping host-seeking behavior (7, 9, 10). While gut microbiota have been well studied, less is known about microbes inhabiting non-gut tissues, including fat bodies and sensory appendages (11, 12). These microbes are ideally positioned to interact with chemosensory organs but remain poorly understood.

Here, we focus on *Escherichia coli*, identified in non-gut tissues of *Aedes albopictus*. Using axenic *Drosophila melanogaster*, targeted reinfection, and co-receptor deletion mutants, we tested whether *E. coli* modulates larval and adult chemosensory behaviors and cuticular hydrocarbon (CHC) profiles. We show that *E. coli* broadly reprograms sensory behaviors, with specific dependence on the ionotropic receptor (IR) co-receptor IR25a. These findings reveal a novel role for *E. coli* in insect chemosensation and suggest microbial cues as potential tools for sustainable insect control.

### E. coli modulates larval sensory-driven behaviors

Insect larvae must integrate short-range sensory inputs—mechanosensation, thermosensation, and phototaxis—to navigate their microhabitats, avoid predation, and secure resources. These modalities are especially critical because larvae, unlike adults, cannot rely on long-distance cues such as odor plumes (1–3).

We observed that tunneling behavior, which reflects substrate exploration and burrowing to avoid desiccation and hypoxia, was reduced in *E. coli*–infected *Canton-S* (wild-type) and *Orco^-^*larvae but remained unchanged in *IR25a^-^*mutants (**Fig. 1A**). Because tunneling normally induces hypoxic stress that shapes larval respiratory physiology (13), these results suggest that *E. coli* influences oxygen-sensing or metabolic regulation in an IR25a-dependent manner. A similar link between microbial metabolites and hypoxia responses has been observed in mosquitoes, where gut microbes shape larval development under variable oxygen conditions (14).

**Figure 1.**
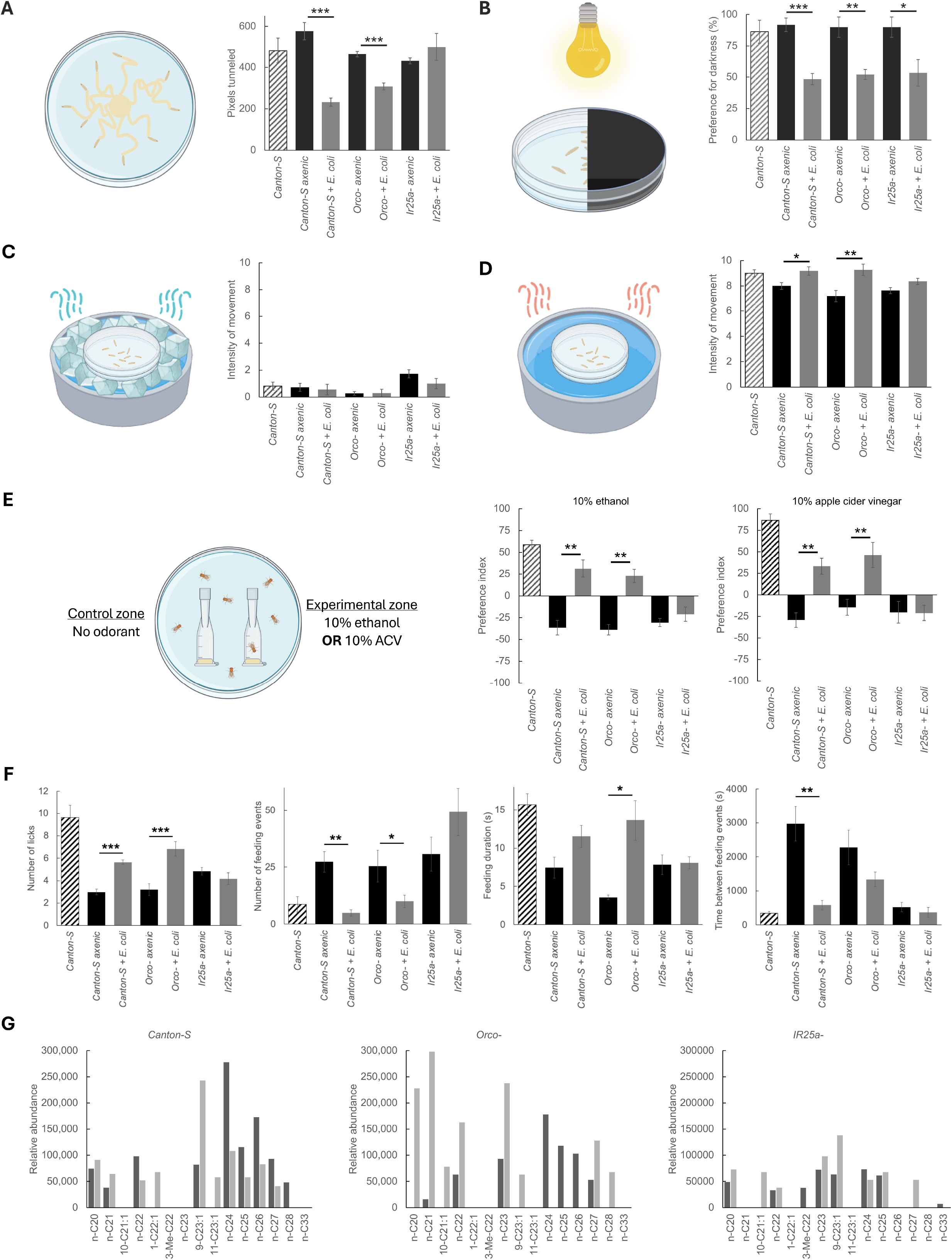
(A-D) Larval behavioral responses in wild-type *Canton-S* and receptor mutant strains under axenic conditions and following *E. coli* reintroduction. (A) Distance tunneled in agar. (B) Preference for darkness. (C) and (D) Intensity of movement in response to heat and cold stimuli, respectively. (E-F) Behavioral assessment of adult wild-type *Canton-S* and receptor mutant strains under axenic conditions and after *E. coli* reintroduction. (E) Olfactory preference measured in a two-choice trap assay, offering flies a choice between standard fly food and food supplemented with 10% ethanol or apple cider vinegar. (F) Gustatory responses to sucrose, including feeding frequency, duration, and latency, enabling analysis of time-dependent changes in preference and individual motivation. *N*=20; *=p<0.05, **=p<0.01, ***=p<0.001. (G) Relative abundance of cuticular hydrocarbons in *Canton-S, Orco^-^*, and *IR25a^-^*strains under axenic conditions (dark grey bars) and following *E. coli* reintroduction (light grey bars). *N*=1.

In phototaxis assays, *E. coli* infection broadly reduced negative phototaxis across all genotypes (*Canton-S, Orco^-^, IR25a^-^*; **Fig. 1B**), indicating a receptor-independent effect. This dampening of light avoidance could arise from systemic immune– neuroendocrine crosstalk, where infection alters neuromodulators that feed back on visual circuits, as described in antibiotic-treated *Drosophila* larvae (15). Similarly, mechanosensory assays revealed heightened withdrawal responses after infection (data not shown), also independent of receptor genotype.

By contrast, thermosensory assays revealed strong IR25a dependence. Wild-type larvae exhibited enhanced responsiveness to heat after infection, whereas *IR25a^-^*mutants failed to show this phenotype (**Fig. 1C–D**). IR25a is a well-established co-receptor for thermal and humidity sensing (16, 17), and our findings suggest that *E. coli* metabolites may act directly or indirectly through these pathways. For larvae, improved thermal discrimination could provide a survival advantage by enabling avoidance of unfavorable microclimates.

Together, these data show that *E. coli* modulates larval behaviors through both receptor-independent pathways (phototaxis, mechanosensation) and IR25a-dependent pathways (tunneling, thermosensation). This dual mode of modulation supports a model in which microbes act not only as environmental signals integrated into sensory circuits but also as systemic modifiers of host physiology.

### E. coli enhances adult olfactory and gustatory behaviors

Adult *Drosophila* rely heavily on olfactory cues to locate food and oviposition sites, often responding to fermentation volatiles such as acetic acid and ethanol (18). In two-choice trap assays, *E. coli*–infected *Canton-S* and *Orco^-^*adults showed increased attraction to apple cider vinegar and ethanol baits, whereas *IR25a^-^*mutants did not (**Fig. 1E**). This indicates that *E. coli* influences adult odor-guided behavior primarily through IR25a-dependent pathways, rather than canonical OR/Orco signaling. Given that IR8a/IR25a complexes mediate detection of acids and fermentation products (19), our results are consistent with microbial metabolites engaging conserved IR circuits.

Feeding behavior assays using the Fly Liquid-Food Interaction Counter (FLIC) revealed that infection enhanced sucrose consumption in *Canton-S* and *Orco^-^*flies but not in *IR25a^-^*mutants (Fig. **1F**). Interestingly, infected flies displayed altered feeding dynamics, including longer but fewer feeding bouts and rapid sequences of consumption, consistent with enhanced motivation to feed. This parallels findings in mammals, where *E. coli* proteins influence appetite and feeding patterns (20). Our data thus provide the first evidence that *E. coli* modulates insect appetite and gustatory responses.

From an ecological perspective, microbial enhancement of attraction to fermentation cues and increased sugar consumption could improve fly foraging efficiency and nutrient uptake, while simultaneously promoting microbial dispersal via contaminated food sources. This is reminiscent of yeast–fly mutualisms, where microbial volatiles attract flies that then vector the microbes to new substrates (21). Here, we extend this concept to bacteria, suggesting that *E. coli* may exploit insect IR pathways to coordinate host foraging and thereby facilitate its own persistence and transmission.

### E. coli alters CHC profiles via IR25a

CHCs are multifunctional traits that protect against desiccation and serve as pheromones for mate recognition and social communication (22). Gas Chromatography-Mass Spectrometry (GC–MS) analyses revealed that *E. coli* infection shifted CHC composition in *Canton-S* and *Orco^-^*flies toward shorter-chain alkanes and alkenes, with a notable increase in 9-tricosene, a male sex pheromone in *D. melanogaster* (**Fig. 1G**). Infected *IR25a^-^*mutants, however, showed no such changes, indicating IR25a-dependent regulation.

These compositional shifts did not alter total CHC abundance, suggesting that microbial influence is targeted to biosynthetic pathways rather than global production. Prior work has shown that microbial colonization can alter CHC profiles by modulating oenocyte transcriptional programs (11, 12). Our findings extend this to *E. coli*, linking bacterial presence to pheromone signaling. Given the role of 9-tricosene in mating dynamics, these microbial effects could reshape reproductive outcomes and potentially promote microbial transmission in microbe-rich environments.

### Mechanistic pathways and ecological consequences

Across both larval and adult stages, *E. coli* exerted IR25a-dependent effects on thermosensation, tunneling, olfactory attraction, and CHC composition, while modulating phototaxis and mechanosensation via IR-independent pathways (**Fig. 2**). This suggests at least two mechanistic routes: (i) direct sensory modulation, where *E. coli* metabolites act as ligands for IR25a-containing receptor complexes, akin to bacterial acid detection in mosquitoes (23); and (ii) systemic modulation, in which infection triggers metabolic or immune changes that secondarily influence sensory neuron responsiveness or peripheral traits such as CHC synthesis.

**Figure 2.**
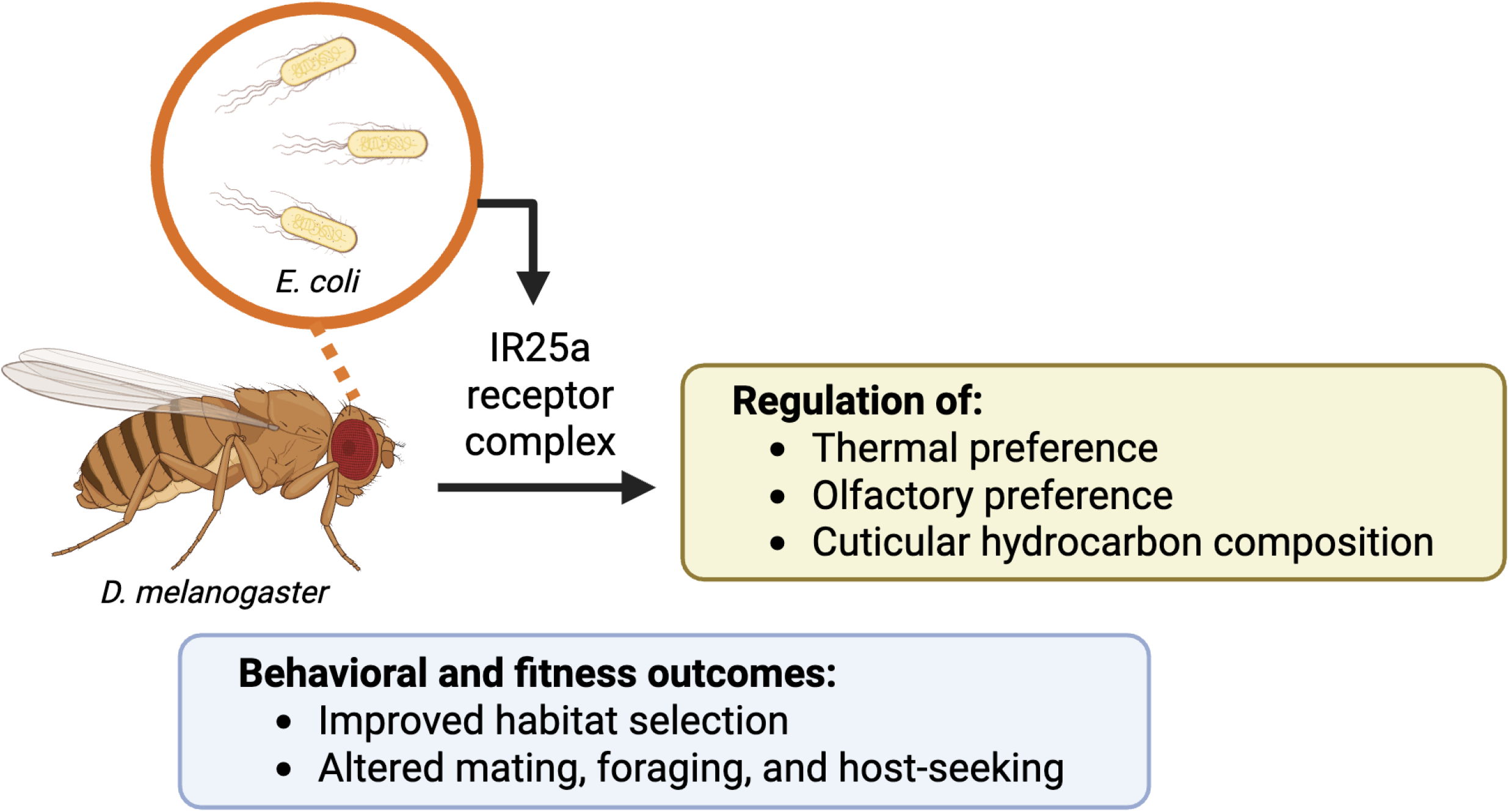
Microbial regulation of insect sensory and behavioral traits. *E. coli* influences multiple *D. melanogaster* sensory modalities through IR25a, leading to benefits in behavior and fitness.

Our findings add to a growing body of evidence that microbes can remodel host sensory landscapes. For instance, *Wolbachia* enhances olfactory receptor expression in *Drosophila* (10), while gut bacteria shape odor-driven feeding preferences (15). The conservation of IR25a across insect orders suggests that microbe-mediated modulation of IR pathways may be a widespread phenomenon.

From an applied perspective, these results highlight microbial cues as potential tools for behavioral pest management. Instead of relying on lethal control strategies, which drive resistance and harm non-target species, microbial metabolites could be deployed to alter insect host-seeking, feeding, or mating behavior in ecologically sustainable ways. Such strategies align with the broader movement toward microbiome-informed interventions in agriculture and vector control.

## Materials and Methods

### Maintaining fly cultures

All experiments used *Drosophila melanogaster* maintained on Nutri-Fly Bloomington Formulation (Genesee Scientific) prepared per manufacturer instructions. Food was autoclaved, cooled to room temperature, then distributed into vials within a biosafety cabinet. Propionic acid (final 0.3% v/v) was added as a fungal inhibitor. Unless otherwise stated, flies were maintained at 25°C, 60% relative humidity (RH), under an 8 h light / 16 h dark (8L:16D) cycle. Populations were age-matched and density-controlled by seeding ∼30 adults per vial for oviposition; adults were removed after 24–48 h to standardize larval density. Experimental cohorts were reared and assayed in parallel to minimize batch and seasonal effects. The lines used in this study were backcrossed to *Canton Special* wild-type strain for ten generations:

*+;IR25a*^-^ *;+* (abbreviated *IR25a*^-^)

*+;+;Orco*^-^ (abbreviated *Orco*^-^)

*Canton Special* (abbreviated *Canton-S*)

*D. melanogaster* are invertebrates and are not subject to IACUC oversight under U.S. federal regulations (Animal Welfare Act, 9 CFR Parts 1–3).

### Preparation of axenic (germ-free) flies

Embryos were collected following Cold Spring Harbor Laboratory protocols. Apple-juice agar plates were prepared and spread with 10 µL yeast paste to stimulate oviposition. 30 adults were placed on each plate and covered with a sterile beaker to form an oviposition chamber, then incubated for 24 h. Embryos were transferred with a moistened paintbrush to 40 µm mesh cell strainers (Olympus). Debris was rinsed away with sterile ddH_2_O. Embryos were dechorionated in 5% (v/v) household bleach for 5 min, then rinsed thoroughly (3× with sterile ddH_2_O). Dechorionated embryos were placed onto fresh, sterile food in autoclaved vials within a biosafety cabinet. Lines were maintained germ-free for 5 generations before use. For verification, spot-plating homogenates of 5 larvae or 5 adults from axenic cohorts on Luria broth (LB) agar (37°C, 24–48 h) to confirm absence of colony growth prior to experiments.

### Reinfection of axenic flies

LB plates were streaked with *E. coli K-12* carrying empty *pET28b* (*ampR*). Isolated colonies were inoculated into LB with ampicillin and grown overnight (16–18 h, 37°C, 200 rpm). For reinfection, 5 mL overnight culture (OD_600_ = 2.0) was aliquoted into 500 µL volumes, pelleted (e.g., 5,000 ×g, 5 min), and each pellet was resuspended in 50 µL LB as concentrated inoculum. This inoculum was spread evenly on the food surface within each sterile fly food vial; a control vial received 50 µL sterile LB only. Axenic adults were flipped into the inoculated vials to oviposit for 7 days, then removed. The resulting progeny (infected generation) were used for assays.

### Environmental control and batch effects

All behavioral assays were performed at 25°C, 60% RH, in complete darkness unless otherwise stated. Biological replicates were run across ≥2 independent days to distribute potential day effects. To minimize seasonal/batch variability, replicates were run within the same week per condition when possible.

### Larval tunneling assay

60 mm × 15 mm Petri dishes with 25 mL 0.5% agar were prepared. A 1 mm diameter central hole was punched and filled with yeast paste. 10 third-instar larvae were briefly rinsed in sterile ddH_2_O with gentle swirling to remove food debris, then placed into the central well. After 30 min, tunneling trails were imaged; distance (pixels) was quantified with identical exposure and magnification across plates in Fiji (RRID:SCR_002285) with a fixed calibration.

### Larval phototaxis assay

60 mm × 15 mm Petri dishes with 25 mL 2% agar were prepared. Half the lid and base were occluded with black paint to create light and dark arenas. 10 third-instar larvae were briefly rinsed in sterile ddH_2_O with gentle swirling to remove food debris, placed in the center of the plate, and exposed to constant overhead illumination for 10 min. Preference (%) = (# in dark / total) × 100%.

### Larval temperature sensitivity assay

10 third-instar larvae were placed in a covered 60 mm × 15 mm Petri dish without agar. For heat, the dish was placed in a water bath held at 32°C; for cold, on ice water; exposure 3 min each. Movement intensity was scored from video (blinded observer) as distance covered per unit time (pixels/min).

### Two-choice trap (olfactory) assay

Two-choice trap assays were designed with vertically oriented traps suspended below the arena to prevent escape. Each baited Eppendorf tube was mounted beneath a small aperture in the arena floor; the aperture was just large enough for a fly to pass, discouraging re-emergence. The vertical orientation ensured once a fly committed, it could not exit easily. 10 virgin adults (5 female, 5 male) were isolated for 2 days post-eclosion and water-starved 24 h, anesthetized briefly on ice, and introduced to the arena for 24 h. Preference index (PI) = ((# in experimental) ™ (# in control)) / (total in traps) × 100%.

### Fly Liquid-Food Interaction Counter (FLIC) assay

To assay single-well food interactions, the FLIC system was configured with the Drosophila Feeding Monitor (DFM) with single chamber tops, connected to the Master Control Unit (MCU). Liquid food (5% sucrose) was loaded into each channel to form a simple voltage divider when contacted by a fly. Signals were logged and translated to volume consumed using a custom R script (RRID:SCR_018386).

### Gas chromatography-mass spectrometry (GC–MS) analysis of cuticular hydrocarbons (CHCs)

CHCs were extracted by hexane wash. 5 adults were placed into 1.5 mL amber GC-MS autosampler vials with 1.0 mL HPLC-grade hexane, vortexed for 1 min. Carrier flow 14 mL/min; set-point pressure 7.05 (poi); oven 40 °C (1 min hold) → ramp 3 °C/min to 300 °C → hold 5 min to elute residuals. Spectra were acquired in scan mode. Relative peak areas were normalized within-sample to total ion current to yield relative abundance profiles.

### Randomization, blinding, inclusion/exclusion criteria

Within each genotype and condition, vials were randomized to assay order using a random number generator. Observers were blinded to genotype and infection status for scoring-based assays (temperature, phototaxis) and for image-based quantification (tunneling). Pre-specified exclusion criteria: vials with visible contamination, assays with escaped or immobile flies (>20% of cohort immobile at start), or hardware/electrical faults (FLIC) were discarded and repeated.

### Data analysis

At *N*=20 plates/vials per group, the study has 80% power (α=0.05, two-sided) to detect standardized effects of ∼0.9 SD in continuous outcomes (or ∼10–12 percentage points for plate-level preference measures, given observed between-plate variability). Analyses were performed at the plate/vial level to avoid pseudoreplication; larvae within a plate are technical subsamples.

Two-tailed Student’s t-tests were used for primary comparisons (infected vs. axenic within genotype). Summary plots show mean ± SEM; significance thresholds were * p<0.05; ** p<0.01; *** p<0.001. Analyses were performed in GraphPad Prism (RRID:SCR_002798) or R Project for Statistical Computing (RRID:SCR_001905).

## Funding

This work is funded by a Rutgers Presidential Postdoctoral Research Fellowship and the Goyette Family Endowment to JSS. The funders had no role in study design, data collection and interpretation, or the decision to submit the work for publication.

## Data Availability Statement

The raw data supporting the conclusions of this article will be made available by the authors on request.

## Acknowledgments

We thank John R Carlson and Lisa S Baik for providing the receptor deletion Drosophila strains. We further acknowledge the research teams of Max M Häggblom and Jeffrey M Boyd for their invaluable comments, and for providing access to materials and instrumentation.

## Conflicts of Interest

The authors declare no conflicts of interest.

## Abbreviations

The following abbreviations are used in this manuscript:

ACV: Apple cider vinegar
CHC: Cuticular hydrocarbon
DMSO: Dimethyl sulfoxide
FLIC: Fly Liquid-Food Interaction Counter
GC-MS: Gas chromatography-mass spectrometry
IR: Ionotropic receptor
LB: Luria broth
OR: Olfactory receptor
PI: Preference index
RH: Relative humidity
SD: Standard deviation
SEM: Standard error of the mean
FLIC: Fly Liquid-Food Interaction Counter
GC-MS: Gas chromatography-mass spectrometry
IR: Ionotropic receptor
LB: Luria broth
OR: Olfactory receptor
PI: Preference index
RH: Relative humidity
SD: Standard deviation
SEM: Standard error of the mean

## Author Bios

### Hazem Al Darwish

#### Department of Biochemistry and Microbiology, Rutgers University, New Brunswick, New Jersey, USA

Hazem Al Darwish is a researcher in the Department of Biochemistry and Microbiology at Rutgers University-New Brunswick. Al Darwish received his B.S. in Biochemistry from Rutgers University, School of Environmental and Biological Sciences (2024). Al Darwish joined the Sun Lab, where his work focuses on the influence of insect-associated microbes on the insect host’s olfactory behavior. He completed an honors thesis as part of the School of Environmental and Biological Sciences’ undergraduate honors program, where he earned the George H. Cook Scholar award.

### Tia Hart

#### Department of Biochemistry and Microbiology, Rutgers University, New Brunswick, New Jersey, USA

Tia Hart is an Honors College undergraduate student at Rutgers University. She is currently pursuing her B.S. in Biological Sciences at the School of Environmental and Biological Sciences. Her experience includes being a Rutgers research assistant for the Aresty program under Dr. Jennifer Sun (2023-present). She studied the influence of microbial symbiosis on insect host behavior. She was also a research assistant for the Rutgers Youth Enjoy Science program under Dr. Moshmi Bhattacharya (2023-2024). In this lab, she studied a potential novel treatment option for Hepatocellular Carcinoma.

Tia plans to put her research assistant experience skills towards pursuing a career in either microbiology or cancer biology research.

## Deep Patel

### Department of Biochemistry and Microbiology, Rutgers University, New Brunswick, New Jersey, USA

Deep Patel is a Rutgers research assistant for the Aresty program under Dr. Jennifer S. Sun’s laboratory at Rutgers University. He is currently pursuing his B.S. in Biotechnology at the School of Environmental and Biological Sciences. Deep plans to apply the skills he gleans from scientific discovery towards a career in pharmaceuticals.

### Sammi Russo

#### Department of Biochemistry and Microbiology, Rutgers University, New Brunswick, New Jersey, USA

Sammi Russo is an undergraduate researcher in Dr. Jennifer S. Sun’s laboratory at Rutgers University. She earned an Associate of Science in Biology from Brookdale Community College (2023) and is currently completing a Bachelor of Science in Microbiology at Rutgers University, expected January 2026. Her research interests focus on the intersection of neurobiology and microbiology. Sammi joined the Sun Lab to study the connections between microbiomes and neurological functions in insects.

### Safiyah Salama

#### Department of Biochemistry and Microbiology, Rutgers University, New Brunswick, New Jersey, USA

Safiyah Salama received her B.S. in Biology and Psychology at the School of Environmental and Biological Sciences, Rutgers University (2025). Her research project was on the influence of *Escherichia coli* on the chemosensory responses in *Drosophila melanogaster* larvae. This project was exploring the role of insect-associated microbes in olfactory circuit shaping. She aimed to know more about insect behavior through her project and the potential it could have in vector control. With a strong interest in biological sciences, she plans to pursue a career as a nurse practitioner. While her research experience was in insect-associated microbes and olfactory circuits, she hopes to bring her scientific expertise to clinical practice, using her analytical mind to patient care and evidence-based medicine.

### Muqaddasa Tariq

#### Department of Biochemistry and Microbiology, Rutgers University, New Brunswick, New Jersey, USA

Muqaddasa Tariq received her B.S. in Biological Sciences and Entomology from the School of Environmental and Biological Sciences at Rutgers University-New Brunswick (2025). She sought research experience with Dr. Jennifer S. Sun in the Department of Biochemistry and Microbiology (2023-2024). During her George H. Cook project, she researched the relationship between environmental factors, mosquito microbiomes, and host chemosensory gene expression. Her long-term career goal is to become an oral surgeon and help people feel more confident with their smiles.

### Jennifer S. Sun

#### Department of Biochemistry and Microbiology, Rutgers University, New Brunswick, New Jersey, USA

Dr. Jennifer S. Sun is an Assistant Professor in the Department of Biochemistry and Microbiology at Rutgers University-New Brunswick. Dr. Sun received her B.A. in Molecular Biology and Biochemistry from Rutgers University (2013) and her Ph.D. in Molecular, Cellular, and Developmental Biology from Yale University (2019). She completed postdoctoral training in Molecular Biology at Princeton University (2019-2021) and Biochemistry and Microbiology at Rutgers University (2022-2024). She was the 2024-2025 President of the Theobald Smith Society: New Jersey Branch of the American Society for Microbiology (ASM). The Sun lab explores the microbiome of insects, focusing on how they shape the host’s olfactory circuits. By studying how microbes change the insect’s sense of smell and resulting behavior (i.e., host-seeking), we can identify novel mechanisms for biting insects at bay. These discoveries will be particularly impactful in developing cost-effective interventions for mitigating insect-borne diseases in low-resource settings.

